# Defining the molecular determinants of titin kinase by analysing distantly evolved fish orthologues

**DOI:** 10.1101/2025.08.04.668337

**Authors:** Till Dorendorf, Peter Gravenhorst, Olga Mayans

## Abstract

Titin kinase (TK) is a pseudokinase specific to the striated muscle of vertebrates. Embedded within the contractile sarcomere and flanked by extensible regulatory tails, TK is thought to sense mechanical signals arising from muscle function. Studies on TK to date have focused narrowly on the human representative. To investigate if a pseudokinase character is a hallmark of TK, we have studied sequences of distantly evolved fish representatives and rationalized conservation patterns by resolving the crystal structure of TK from medaka isoform *b.* We find that sequence alterations in catalytic motifs involved in ATP and magnesium binding, respectively θxK (θ: bulky hydrophobic residue) and EFG, are evolutionarily conserved in TK. Beyond the kinase domain, N- and C-terminal flanking tails show remarkable structural similarity across human and medaka orthologues, even though sequence conservation is limited to individual residues and short motifs: a YD-motif in the N-terminal tail; a [R/K]H[R/K]RYY sequence, a R-7x-R motif and position −2 of the latter in the C-terminal tail. We observe that motifs in the C-terminal tail consistently covary with the divergent functional motifs of TK, being part of its pseudokinase signature. In contrast, the proposed inhibition of the catalytic aspartate by a tyrosine residue from the P+1 loop is not significantly conserved outside mammals. Finally, based on these features and sequence clustering analysis, we propose a classification of titin gene duplicates from fish into *a* and *b* isoforms (*ttna* and *ttnb*) that can assist future comparative studies. A curated genomic annotation is provided here.

## Introduction

The giant protein titin (∼3-4MDa) plays a central role in generating, sensing and transmitting passive force in the muscle sarcomere. In addition to being an important regulator of sarcomere mechanical performance, titin has mechanosensory functions that are central to triggering the adaptive response of the myofibril to mechanical demand (Krüger & Kötter, 2016; Linke, 2023). The delivery of these functions is enabled by the architecture of titin, a multi-domain elastic protein that spans the half-length of the sarcomere, from Z-disc to M-line (more than 1 µm in length). Titin is primarily composed of small immunoglobulin (Ig) and fibronectin type III (FnIII) domains, which jointly account for over 90% of its mass (Bang et al., 2001). A single signalling domain is found in titin, a kinase domain (titin kinase; TK) near the C-terminus of the protein, located in the sarcomeric M-line. The function of TK has remained enigmatic ever since its discovery over 30 years ago (Labeit & Kolmerer, 1995). However, being embedded within the contractile apparatus of the sarcomere, it is regarded as a prime candidate to act as a mechanoreceptor in the sensing and transduction of mechanical signals arising from muscle function.

TK mechanosensing is thought to be mediated by extensible, regulatory tail sequences that flank N- and C-terminally the kinase domain (termed, N-terminal linker: NL; C-terminal regulatory domain: CRD). Structural characterization of human TK showed that the NL and CRD flanking tails bind tightly to the interlobular kinase hinge region and active site, respectively (Mayans et al., 1998; Bogomolovas et al., 2021). Structural studies of the close homologue twitchin kinase, both from *Caenorhabditis elegans* (*ce*TwcK) (Hu et al., 1994; Kobe et al., 1996; von Castelmur et al., 2012) and *Aplysia californica* (*ac*TwcK) (Kobe et al., 1996) (40% and 41% sequence identity to human TK, respectively) revealed a closely similar architecture. For *ce*TwcK, the tails have been shown to completely inhibit phosphotransfer catalysis (Kobe et al., 1996; von Castelmur et al., 2012), so that kinase activation requires the removal of the regulatory tails from their observed position. It is postulated that in sarcomeric kinases activation is elicited by mechanical force, where the pulling forces that develop in the sarcomere during muscle function cause the tails to unravel, thereby exposing the kinase active site (Gräter et al., 2005; Puchner et al., 2008; von Castelmur et al., 2012; Bogomolovas et al., 2021). Support to the physiological relevance of this mechanism was brought by a recent study that monitored fluorescence resonance energy transfer (FRET) in live, freely swimming *C. elegans* worms to reveal conformational changes in *ce*TwcK during muscle activity *in vivo* (Porto et al., 2021).

While tail removal can be expected to lead to kinase activation triggering a force-dependent signalling pathway in the cell, it is the case that TK is a pseudokinase with questionable phosphotransfer delivery (Bogomolovas et al., 2014). In TK, two catalytic motifs are degenerated when compared to canonical kinases: MAK (instead of the canonical AxK sequence) in the ATP-binding pocket and an EFG motif (instead of the conserved DFG sequence) in the magnesium binding site (Manning et al., 2002; Bogomolovas et al., 2014). The AxK motif in strand-β3 of the kinase fold contains the essential catalytic lysine that binds and positions the non-transferable α- and β-phosphates of ATP. In active kinases, small hydrophobic residues occupy position 1 of this motif (typically alanine or valine), as this position forms the bottom of the cavity that accommodates the nitrogenated base in ATP (Endicott et al., 2012). In TK, the bulkier methionine residue at this position is speculated to sterically conflict ATP binding (Bogomolovas et al., 2014). In the DFG motif, aspartate chelates the magnesium ion required for catalysis and is relevant for γ-phosphate transfer. Its substitution for glutamate drastically reduces phosphotransfer activity in canonical kinases (Schu et al., 1993; Brown et al., 1995; Bogomolovas et al., 2014). Thus, it is expected that also in TK the presence of a glutamate troubles activity. In agreement with this view, validated substrates and phosphotransfer-dependent pathways of TK are yet to be identified.

In contrast, the scaffolding functions of TK are better understood. TK associates with the E3 ubiquitin ligases MURF1 (Bogomolovas et al., 2014; Bogomolovas et al., 2021) and MuRF2 (Lange et al., 2005) and the autophagy receptors p62 and Nbr1 that promote the selective autophagy of poly-ubiquitinated proteins (Lange et al., 2005; Bogomolovas et al., 2021). While the functional outcomes of such scaffolding are not yet fully clarified, it has been proposed that the release of MuRF2 from its TK locus in inactive muscle cells leads to the suppression of the nuclear serum response factor, which in turn hinders the transcription of anabolic genes as corresponding to mechanical inactivity (Lange et al., 2005). A more recent study (Bogomolovas et al., 2021) found that ubiquitination of the TK locus by MuRF1 led to targeting by Nbr1 and p62 that associate with the ubiquitin moiety, possibly being functionally related to sarcomere loss processes. It was suggested that the extension of the kinase tails, resulting from muscle activity, decreased ubiquitination levels of the TK locus and, thereby, reduced the Nbr1 and p62 targeting of TK, thereby correlating muscle activity with sarcomere preservation. Taken together, these findings have led to regard TK as a scaffold based signalling hub for mechanosensing with links to the nucleus and protein turn-over pathways, regulated by the conformational state of TK.

Notably, studies of TK to this date have focused narrowly on the human representative, so that it is unclear to which extent the properties of TK are conserved across vertebrates. In order to identify the molecular determinants of TK, in this work we have studied sequence conservation in distantly related TK representatives from fish, the most ancestral class of vertebrate. Fish constitute a rich source of diversified protein sequences, having evolved over long evolutionary distances and containing duplicate gene variants dating back to the teleost-genome duplication (TGD) that occurred 267-286 million years ago (Qi et al., 2024). Zebrafish (*Danio rerio*) and medaka (*Oryzias latipes*) are two teleostei model fish organisms that represent two modern phylogenetic groups. Medaka belongs to the more recently emerged monophyletic clade of the neoteleostei, whereas the zebrafish branched off earlier in the teleost evolution. Among fish, the titin copies of the zebrafish are the best studied so far. These two titin proteins (ttna and ttnb isoforms) have been ascribed different functions. Isoform ttna is highly expressed in cardiac muscle (ratio of 2:1 ttna:ttnb) and is essential for sarcomere formation in the heart, whereas isoform ttnb is dispensable in this muscle type (Seeley et al., 2007). In contrast, the expression ratios of ttna and ttnb in skeletal muscle sarcomeres are similar and both isoforms are required for sarcomere formation (Seeley et al., 2007; Zou et al., 2015). Similar to zebrafish, medaka also contains two copies of the titin gene, but these are largely uncharacterized. Guided by these model systems, we have performed a sequence analysis of titin isoforms and further compared results to data from mammalian representatives. Complementing this analysis, we have elucidated the crystal structure of TK from the ttnb isoform of *Oryzias latipes* (medTK*b*) that allows us to rationalize sequence conservation patterns. This analysis has revealed the molecular determinants of TK and facilitates the understanding of titin-related sarcomeric kinases.

## Methods

### Cloning

The coding DNA for the TK domain of ttnb from *Oryzias latipes*, medTK*b*, (residues 27887–28241; NCBI XP_023806503) was synthesised commercially (Biocat, DE). The gene was cloned into the vector pETtrx1a (J. Bogomolovas) using NcoI and KpnI restriction sites, as described previously for human TK (Bogomolovas et al., 2022). This vector fuses a His_6_-tagged thioredoxin and a tobacco etch virus (TEV) protease cleavage site N-terminally to the gene insert of interest. The construct was confirmed by sequencing (Eurofins, DE).

To ease structural annotation in this work and facilitate comparison to previously reported crystal structures of human TK, residue M27887 in entry XP_023806503 is set as residue 1 in our study.

### Protein Production

The medTK*b* kinase was expressed recombinantly in *E. coli* SoluBL21 (Amsbio, UK) in TB medium supplemented with 50 µg/mL kanamycin, as described for human TK (Bogolomovas et al., 2022). Cultures were grown at 37 °C to an OD_600_ ≥ 1. Upon cooling to 18 °C, expression was induced with 0.5 mM IPTG and growth continued for a further 18 h. Cells were harvested by centrifugation. The pellet was resuspended in lysis buffer containing 20 mM HEPES pH 8.0, 250 mM NaCl, 5 mM imidazole, 0.2% Triton-X-100, 1 mM dithiothreitol (DTT) and supplemented with cOmplete^TM^ ULTRA EDTA-free protease inhibitors (Roche, CH). Cell lysis was done by sonication and the lysate subsequently clarified by centrifugation. The supernatant was applied to a Ni^2+^ HisTrap HP column (Cytiva, SE) pre-equilibrated in lysis buffer and proteins eluted with 300 mM imidazole in lysis buffer without detergent. Tag removal was by incubation with TEV protease in 25 mM Tris pH 8.0, 50 mM NaCl, 1 mM DTT, overnight at 4 °C. Proteins were further purified using ion exchange chromatography on a HiTrap SP column (Cytiva, SE) and gel filtration on a Sephadex S200 16/60 column (GE Healthcare, DE) equilibrated in 25 mM Tris-HCl pH 8.0, 50 mM NaCl, 1 mM DTT. Samples were stored at 4°C until further use.

### X-ray crystallography

The medTK*b* sample was concentrated to 11.6 mg/mL and used in crystallization trials that used the sitting drop vapour diffusion method in ARI Intelliplates (Art Robbins Instruments, USA) incubated at 18 °C. Drop composition was a mixture of 100 nl protein : 100 nl reservoir solutions. Crystals grew in 20% [w/v] PEG 8000, 100 mM HEPES pH 7.5. For X-ray data collection, the crystals were harvested and flash-frozen in liquid nitrogen using mother liquor supplemented with 30% [v/v] ethylene glycol as cryo-protectant. X-ray diffraction data were collected at beamline P14 (EMBL, DESY, Hamburg, DE) under cryo-conditions (100 K). For data processing, the XDS/XSCALE suite was used (Kabsch, 2010). Phasing was done by molecular replacement in PHASER (McCoy et al., 2007) using human TK as template (PDB code 4JNW). An initial atomic model was built automatically using Arp/wArp (Langer et al., 2008) and further manual building was performed in COOT (Emsley et al., 2010). Model refinement was carried out in Phenix.refine (Liebschner et al., 2019) applying TLS refinement and non-crystallographic symmetry (NCS), with each chain being a TLS and NCS group. X-ray data statistics and model parameters are given in **Table 1**. The first eleven residues of chain A and the first nine of chain B (of which the first two residues originate from the vector) are not visible in electron density maps, as a result the first residues contained in the crystal structures are L10 and K8 for chains A and B, respectively (residue L27896 and K27894 in NCBI XP_023806503).

**Table 1:** X-ray data statistics and model parameters.

|  | **medTKb** |
| --- | --- |
| PDB code | 9QJ2 |
| Space group | P2<sub>1</sub> |
| Cell dimensions: a,b,c (Å) | 61.74, 56.08, 120.25 |
| Cell angle: $\alpha$, $\beta$, $\gamma$ (°) | 90.00, 104.28, 90.00 |
| Copies in ASU | 2 |
| **Data Processing** |  |
| Beamline | DESY beamline P14 |
| Detector | EIGER2 16M |
| Wavelength (Å) | 0.9763 |
| Resolution (Å) | 30-2.1 (2.15-2.1)<sup>a</sup> |
| No. Reflections | 46545 (3175)<sup>a</sup> |
| R<sub>sym</sub>(I) (%) | 12.2 (258.2)<sup>a</sup> |
| $\langle I/\sigma(I) \rangle$ | 10.55 (0.91)<sup>a</sup> |
| CC1/2 (%) | 98.8 (37.0)<sup>a</sup> |
| Completeness (%) | 99.1 (99.2)<sup>a</sup> |
| Multiplicity | 6.88 (7.04)<sup>a</sup> |
| **Model Refinement** |  |
| No. working/free reflections | 46535 / 1427 |
| Rwork/Rfree (%) | 19.02 / 22.72 |
| No. residues / atoms protein | 688 / 5516 |
| NCS R.m.s.d (Å) | 0.284 |
| No. Atoms Solvent | 215 + 11 x EDO<sup>b</sup> |
| R.m.s.d. bond length (Å) | 0.007 |
| R.m.s.d. angles (°) | 0.900 |
| Ramachandran plot |  |
| Favored/disallowed (%) | 98.68/0.0 |

<sup>a</sup> number in parentheses are the values for highest resolution shell;
<sup>b</sup> EDO: 1,2-ethanediol.

### Sequence comparison

The sequence of human TK, including its NL and CRD segments, (UniportKB: Q8WZ42, residues 32150-32491) was used as query in a blast search of related sequences (https://blast.ncbi.nlm.nih.gov/Blast.cgi) that yielded 5034 entries. The sequences were curated to include proteins from vertebrates only (i.e. excluding potentially mislabeled titin-like kinases from invertebrates). In addition, entries labelled as myosin light chain kinase or annotated as LOW_QUALITY were excluded. Finally, the sequences were made unique to exclude redundancy deriving from different titin isoforms with sequence variations in regions other than the kinase domain. This left 738 sequences including the clades of reptiles, fish, amphibians and mammals; 459 of those were sequences from fish and mammals, of which 320 sequences belonged to fish.

The 320 sequences of fish were then clustered using PaSiMap (Su et al., 2022) implemented in JalView (Morell et al., 2024). Cluster plots were generated with RStudio (https://posit.co/download/rstudio-desktop). As this revealed that fish sequences segregated into different clusters, a next analysis looked at the clustering of all 459 sequences of fish and mammals to explore if mammalian TK is similar to one of the fish groups or forms an own cluster.

Finally, in order to quantitate the natural residue variability at functional loci of the kinase fold, the 459 sequences were aligned with ClustalO (https://www.ebi.ac.uk/jdispatcher/msa/clustalo?stype=protein). Partial sequences that did not cover the full kinase region were removed (leaving 452 sequences) and then residue type per position within the multiple sequence alignment quantified and illustrated using pie-charts.

## Results

### Comparing Titin Kinases from fish reveals evolution of distinct isoforms

In order to undertake a comparative study, TK sequences from fish were collected by performing a similarity search in Blast (https://blast.ncbi.nlm.nih.gov) using the sequence of human TK as a query. Upon curation, the search yielded 320 sequences from teleost fish (provided in **Supp Mat Table S1**). Aiming to reveal sequence subgroups sharing closer similarity within the set, the sequences were subjected to clustering analysis using PaSiMap (Su et al., 2022). PaSiMap is a sequence clustering algorithm that transforms pairwise similarity values into correlation coefficients and projects them into a multi-dimensional space (Su et al., 2022; Morell et al, 2024). Each sequence is graphically depicted as a vector in space from the coordinate origin. PaSiMap analysis caused the fish TK sequences to segregate into four clusters (the interactive 3D-vector map from PaSiMap is provided as **Supp Mat S2**). A 2D-representation of dimension 2 vs dimension 3 of the PaSiMap vector map (**Figure 1A)** shows each sequence cluster localizing to a quadrant of the coordinate system. Dimension 2 separates the sequences according to TK isoforms, *a* and *b*, where TK copies that emerged through gene duplication in a same organism localize to different clusters. The TK domain of ttna from zebrafish localizes to the cluster located below the axis origin (0), whereas TK from zebrafish ttnb is found in the cluster above 0. Using as reference these annotated titin isoforms from zebrafish, we extend here the *a* and *b* nomenclature to the rest of TK sequences in the corresponding clusters. Aiming to reveal whether this classification of TK sequences is representative of full-length titin proteins, we performed an equivalent PaSiMap cluster analysis on selected full-length titin sequences from fish. Specifically, the analysis included both titin isoforms from medaka, ocellaris clownfish [*Amphiprion ocellaris*] and tuna [*Thunnus albacares*] from the neoteleostei clad and zebrafish [*Danio rerio*], goldfish [*Carassius auratus*] and yellowhead catfish [*Tachysurus fulvidraco*] as non-neoteleostei (**Supp Fig S1**). This analysis reproduced the results obtained for the isolated TK domains, leading us to conclude that the TK domain is a valid proxy for the classification of full-length titin genes in fish. Thus, we propose here extending the nomenclature ttna and ttnb to the full-length titin protein for the other 318 fish species studied in this work. The new gene annotation is listed in **Supp Mat Table S1**.

**Figure 1:**
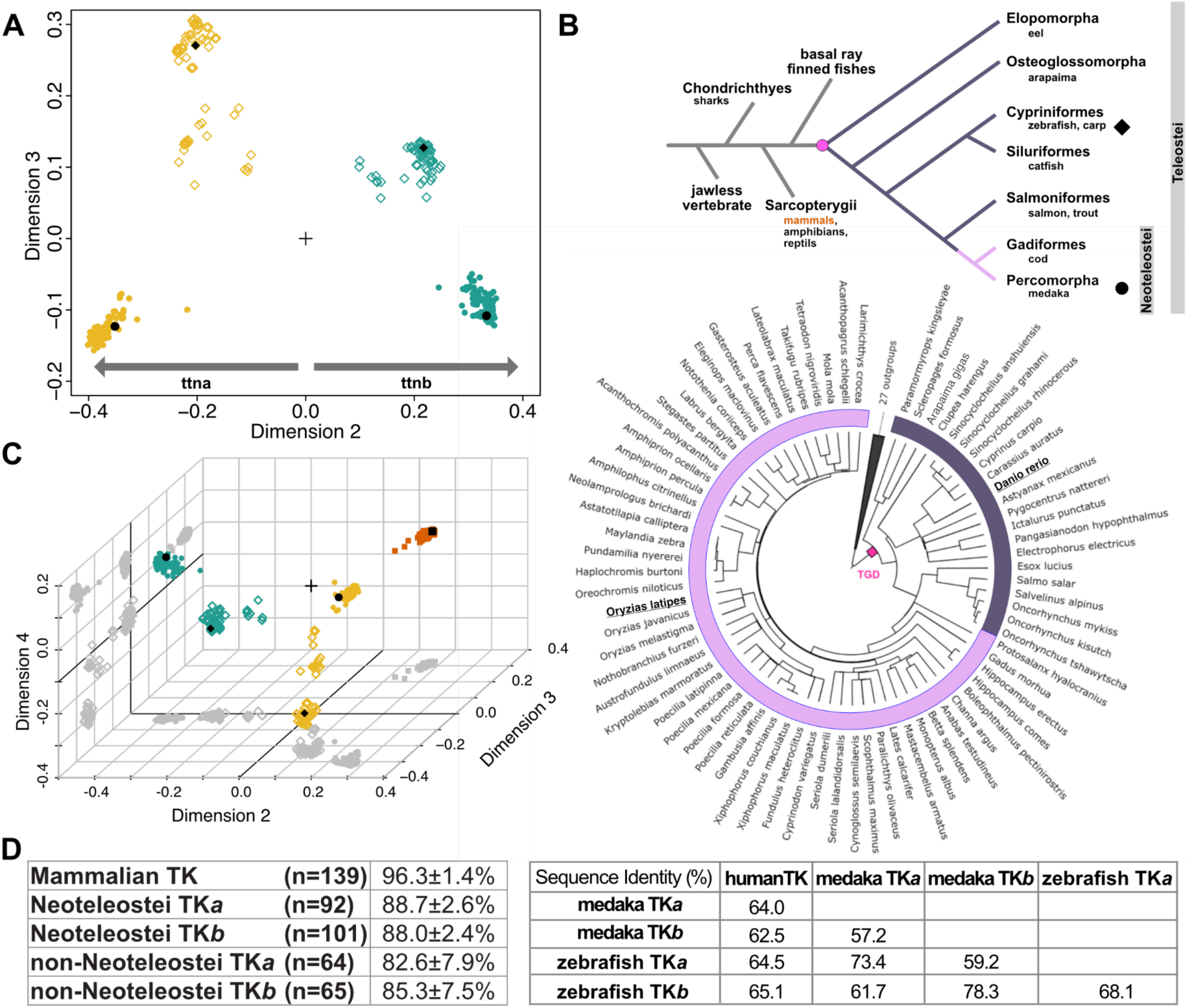
PaSiMap based comparison of Titin Kinase sequences. **A.** Clustering of TK sequences from fish as visualized in a 2D-vector map from PaSiMap (Su et al., 2022). The sequences encompass the NL segment, the kinase domain and the CRD-tail of TK. Four sequence clusters are identified. Yellow and teal colours indicate TK isoforms *a* and *b*, respectively. Fish groups are indicated by diamond and circular data points, where diamonds are used for non-neoteleostei and circles indicate neoteleostei. TK sequences from model organisms representative of each group, zebrafish (*Danio rerio*; ◆) and medaka (*Oryzias latipes*; □), are indicated with black symbols; **B.** A simplified representation (*upper panel*) of vertebrate evolution including the branching of the mammalian lineage and a basic phylogenetic tree of fish. A more exact fish phylogenetic representation (*lower panel*) is also shown (adapted from Parey et al., 2022). In both representations, the point of teleost-genome-duplication (TGD) is indicated as a dot in magenta, with neoteleostei being shown in pink and older fish in aubergine; **C.** PaSiMap clustering of fish and mammalian TK sequences. Fish colouring is as in A., mammalian TKs are shown as brown squares (human TK is highlighted as a black square; ▪). To aid 3D-visualisation, data point projections on the axial planes are shown as shadows in grey; **D.** Matrix of sequence identities (expressed in percentage) within PaSiMap sequence clusters corresponding to animal groups (*left*) and across protein representatives of main groups (*right*). Within a same cluster, sequence identity is calculated as average sequence identity ± standard deviation; number of sequences in each group is stated.

In the PaSiMap vector map, sequence segregation in dimension 3 correlates with fish group (**Figure 1B**); namely, all sequences from neoteleostei (*e.g.* medaka [*Oryzias latipes*], ocellaris clownfish [*Amphiprion ocellaris*] or tuna [*Thunnus albacares*]) have values below 0, while sequences from evolutionarily older fish (*e.g.* zebrafish [*Danio rerio*], goldfish [*Carassius auratus*] or yellowhead catfish [*Tachysurus fulvidraco*]) have values above 0. This indicates that these fish groups can also be identified and catalogued through the analysis of TK.

Importantly, PaSiMap places isoform type in dimension 2 and fish group in dimension 3. As the order of the dimensions in the clustering analysis corresponds to the prominence of the sequence divergences, this signifies that differences between TK isoforms in a same organism are larger than differences in the same isoform across fish groups. This deduction reflects excellently the order of evolutionary events in fish, and therefore, the evolutionary distances of the sequences studied (**Figure 1B**).

In order to investigate how the diversified TK*a* and TK*b* domains from fish relate to the better characterized human TK, we extended the PaSiMap analysis to mammalian sequences. The original Blast search described above also yielded 139 sequences of mammalian TKs, which were then added to the fish counterparts and subjected to a new PaSiMap clustering. The resulting PaSiMap vector map (**Figure 1C**; the interactive PaSiMap 3D-vector map is provided as **Supp Mat S3**) segregated mammalian TK sequences into an additional fifth cluster, suggesting that fish TK*a*, TK*b* and mammalian TK form distinct groups without mutual overlap. To analyse the relation of sequences within individual clusters and across clusters, the inter- and intra-cluster sequence identity levels were quantitated (**Figure 1D**). Within clusters, sequence identity is high in all cases, but mammalian TK sequences share the highest identity levels (96.3%) even though the sequences originate from phylogenetically distant animals such as the egg-laying mammal *Tachyglossus aculeatus*, the marsupial *Monodelphis domestica* or the narwhal. Fish TK sequences also show high conservation levels within the clusters, but the older non-neoteleostei group is more diversified, as it could be expected. To exemplify sequence identity across clusters, we compared human, medaka and zebrafish TK representatives (**Figure 1D, right**). This analysis yielded values ranging from 57.2% to 78.3% and confirmed that conservation is higher within each isoform type, i.e. isoforms *a* share higher identity to each other across all fish groups than to isoforms *b* within the same organism, which is also true for isoforms *b.* This agrees excellently with deductions drawn from examining PaSiMap dimensionality (above).

### Structural characterisation of medaka TK*b* reveals a conserved fold

To gain a better understanding of the molecular diversity of TK kinases, we have elucidated the crystal structure of the most divergent representative from a fish model organism, TK isoform *b* from medaka (medTK*b*), to 2.1Åresolution (**Table 1**; **Figure 2A,B**). The construct comprised the NL-kinase-CRD regions. The crystal form obtained in this study contained two molecular copies in the asymmetric unit, which were essentially identical to each other (RMSD_Cα_ = 0.284 Å for all protein residues). The crystal structure of medTK*b* closely resembles that of human TK (Mayans et al., 1998; Bogomolovas et al., 2014), with which medTK*b* shares 62.5% sequence identity (**Figure 2C**). Remarkably, the highly conserved 3D-fold of TK extends to its NL and CRD flanking extensions even though sequence conservation within these tails is low and mostly confined to specific single residues or very short motifs (described below). Only a minor deviation of the fold can be observed, localized to the 8-residue loop region between helices αR1 and αR2 in the CRD tail. In conclusion, the overall topology of medTK*b* is closely coincident with that of the well-studied human orthologue.

**Figure 2:**
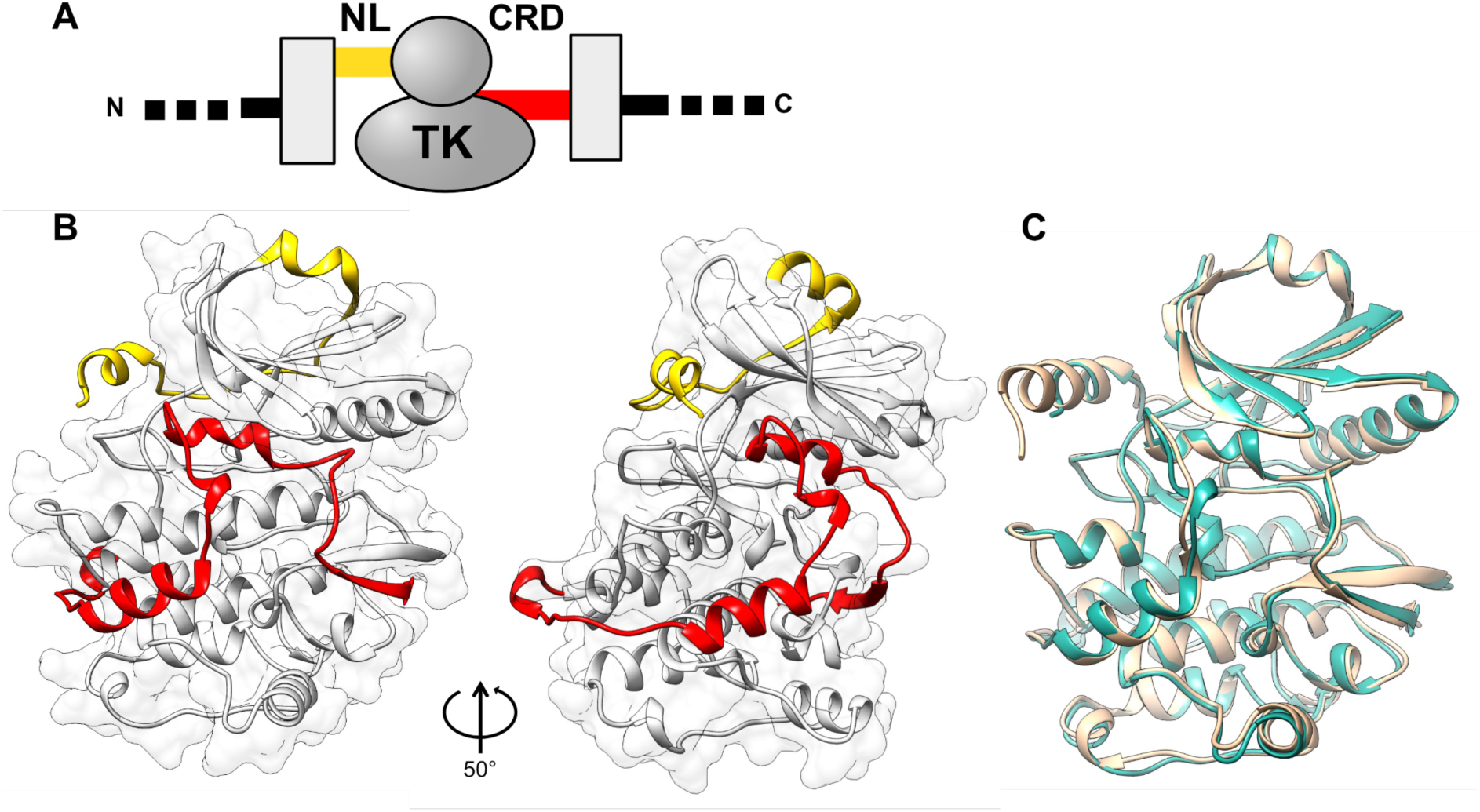
Crystal structure of medaka TK*b*. **A.** Schematic depiction of the domain composition of the kinase region of titin proteins. The kinase domain (TK) is flanked N-terminally by a linker sequence (NL, yellow) that joins it to the preceding FnIII-domain within the full-length titin protein. C-terminally, TK is flanked by a regulatory tail extension (CRD, red), which is followed by an Ig domain in the titin chain. The neighbouring FnIII and Ig domains are shown here as grey boxes; **B.** Crystal structure of medaka TK*b* shown in two orientations, corresponding to a rotation of ∼50° around an axis contained within the plane of the image. Colour code is as in A. The surface outline corresponds to the kinase domain (grey); **C.** Superimposition of medaka TK*b* (teal) with the corresponding segment of human TK (extracted from the larger multi-domain structure in PDB entry 6YGN, gold); (RMSD_Cα_ = 1.10 Å for all 345 overlapping C_α_-atoms; calculated using UCSD Chimera; https://www.cgl.ucsf.edu/chimera).

### Conservation of pseudokinase features in the TK domain

Catalytic readiness in kinases is achieved through a finely choreographed set of conformational changes that collectively make the kinase transition from an “off” to an “on” state (Taylor & Kornev, 2011; Kornev & Taylor, 2015). The catalytically ready state is defined by: *i)* the position of helix αC, which allows the formation of a salt bridge between its strictly conserved glutamate and the invariant lysine from strand β3 (in the conserved motif AxK) that coordinates and productively orients ATP for catalysis; *ii)* an open conformation of the activation loop that allows access to the active site; *iii)* the assembly of two spines (R[regulatory]-spine and C[catalytic]-spine) consisting of stacked hydrophobic residues that promote the mutual rearrangement of the kinase lobes; *iv)* the “in” arrangement of a conserved DFG motif, where the F residue is oriented towards the interior of the kinase completing the R-spine and the D residue, involved in the coordination of the Mg^2+^ cofactor, is accessible in the active cleft. While these conformational changes in kinases are commonly induced by the binding of ATP, cytoskeletal and sarcomeric kinases of the DMT clad appear to pre-exist in a catalytically primed state regardless of ATP binding (Temmerman et al., 2013). With the exception of *i),* these hallmarks of the active kinase state are displayed by the autoinhibited human TK (Mayans et al., 1998; Bogomolovas et al., 2014) as well as medTK*b* in this study, confirming that TK domains pre-exist in a semi constitutively active state, uniquely blocked from activity by the flanking tails (**Figure 3A,B**). In both medTK*b* and human TK, the formation of the essential salt bridge between the invariant lysine in strand β3 (K68 in medTK*b*) and the conserved glutamate residue (E83) from helix ⍺C is prevented by the binding of the CRD tail into the ATP pocket, which displaces helix ⍺C into a tilted “open” conformation. In medTK*b,* this displacement results in a distance of 4.92±0.17 Åbetween the amino group of K68 and and the oxygen atoms of the carboxyl group of E83. Interatomic distances are also over 4.9 Å in human TK. To date, no crystal structure of an uninhibited TK domain (or a close TwcK homologue) is available. This, due to the difficulty of producing viable recombinant samples in the absence of the flanking tails, which contribute significantly to kinase stability (Bogomolovas et al., 2022).

**Figure 3:**
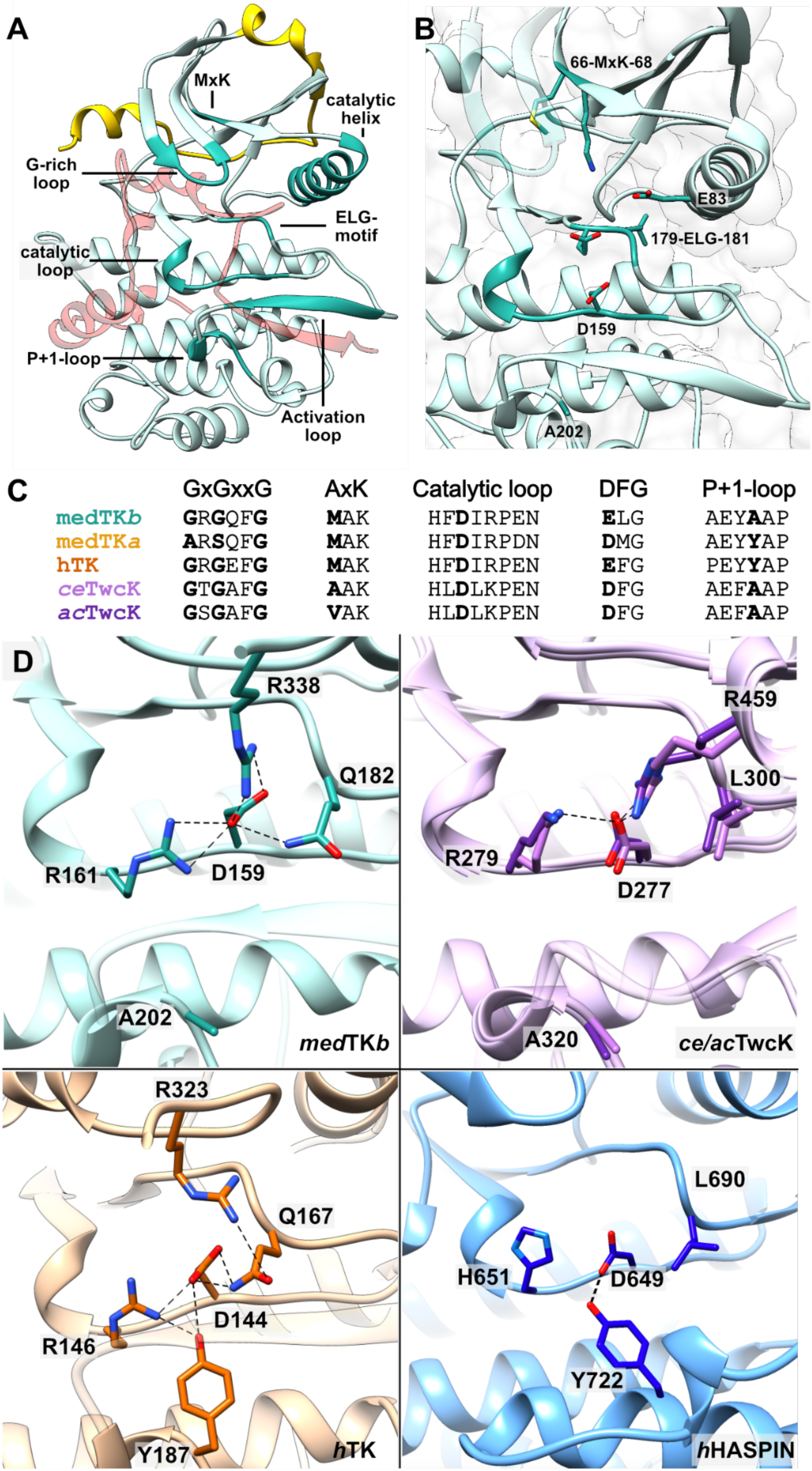
Catalytic features of medTK*b*. **A.** Crystal structure of medTK*b*, where functionally relevant features are highlighted in turquoise and annotated; namely, the ATP-binding Gly-rich loop, the MAK motif in strand β3, the metal cofactor chelating ELG motif, the activation loop, the P+1 loop that harbours the putatively inhibitory tyrosine in human TK, and the catalytic loop that harbours the catalytic aspartate. The NL linker is shown in yellow and the CRD tail in transparent red; **B.** Active site of medTK*b*; **C.** Sequence alignment of selected functional motifs in TK from medaka isoforms, human TK, and TwcK of known 3D-structure; **D.** Comparative overview of the interactions sustained by the catalytic aspartate in representative kinases, where the structures shown are medTK*b* (this study), *C. elegans (ce;* lilac) and *A. californica (ac;* purple) TwcK (PDB: 3UTO and 1KOB, respectively) (residue numbering shown belongs to *ce*TwcK), human TK (PDB: 4JNW), and human HASPIN kinase (PDB: 2WB8). For visual clarity, the CRD is excluded from the images of TK and TwcK structures. Hydrogen bonds are shown as dotted lines.

In medTK*b*, the two sequence deviations in the canonical DFG and AxK motifs that led to question the phosphotransfer activity of mammalian TKs are also present, here being <u>E</u>LG and <u>M</u>AK (**Figure 3B,C**). The ELG sequence of medTK*b* is also in the “in” conformation (**Figure 3B**). The substitution of phenylalanine to leucine in this motif does not appear to have structural consequences and it still results in the effective formation of the hydrophobic R-spine (residues L87, L98, H157, L180). The second hydrophobic spine (C-spine) in canonical kinases is completed by the binding of ATP via the purine base moiety. We observe that in medTK*b*, the C-spine (residues V53, M66, I120, I165, V166, Y167, L224, L228) is completed by residue A328 from helix ⍺R2 of the CRD that packs against the MAK sequence (see further description below). A comparative exploration reveals that also in the autoinhibited states of human TK and TwcK from both *C. elegans* and *Aplysia*, the C-spine is completed by a residue from helix ⍺R2 of the CRD that mimics ATP binding, packing into the ATP binding pocket and against the AxK sequence of strand β3. For this packing to occur efficiently in TKs, the residue from CRD-helix ⍺R2 must necessarily be small. It is unclear whether the small alanine residue from the CRD in TK can be efficiently replaced by the bulkier nitrogenated base of the ATP. The residue covariation in strand β3 and helix aR2 in TKs appears to support the view that the AtoM residue substitution in the AxK motif conflicts ATP binding. In brief, medTK*b* shares many features with DMT kinases structurally characterised to date, but holds the same deviating features of human TK, thereby, also retaining the pseudokinase character (**Figure 3C**).

A further feature of human TK that has complicated the exploration of its potential phosphotransfer activity is the speculated inhibition of the catalytic aspartate (D144; numbering as in PDB: 4JNW) by a tyrosine (Y187) from the P+1 loop to which it hydrogen bonds (**Figure 3C,D**) (Mayans et al., 1998). Both D144 and Y187 residues are embedded within a hydrogen bond interaction relay network with residues R146, Q167 and R323. It is thought that the inhibition might be relieved by the phosphorylation of Y170 (Mayans et al., 1998; Lange et al., 2005) in a similar fashion as observed in kinases ERK2 (Mace et al., 2013) and IRK (Wu et al., 2008). However, the presence of tyrosine at this position is not necessarily inhibitory as shown by the atypical, but active kinase HASPIN, which has a tyrosine in the P+1 loop that interacts with the catalytic aspartate (**Figure 3D**) and does not require of phosphorylation for activation (Eswaran et al., 2009). Thus, the inhibitory character of the tyrosine in human TK remains to be established. While the tyrosine in the P+1 loop is also present in the sequence of medTK*a* (**Figure 3C**), where a predicted 3D-model suggests that it also interacts with the catalytic aspartate (**Figure S2**), the residue is not conserved in medTK*b*, which contains instead an alanine residue (A202) at that position (**Figure 3C,D**). We conclude that this tyrosine does not constitute a general regulatory mechanism in TKs.

### Conservation in the N-terminal linker is limited to a YD kinase-binding motif

In titin proteins, the TK domain is always flanked N-terminally by a linker sequence (NL), approx. 33 residues in length, that joins it to the preceding FnIII-domain within the chain. Even though its sequence composition is highly variable, the crystal structure of medTK*b* reveals that the NL tail is a structurally conserved feature in TK, consisting of a short N-terminal α-helix followed by an extended chain that wraps around the N-terminal kinase lobe (**Figure 2A & 4A,B**). Sequence conservation in the NL tail is confined to a short, strictly conserved YD motif, commonly preceded by an asparagine residue (73.8% of all TK sequences). Structurally, the motif interacts with the inter-lobular hinge-region of the kinase domain, being the main responsible for the packing of the NL sequence against the kinase. In medTK*b*, the hydroxyl-group of residue Y12 forms a hydrogen-bond to the side chain of E43 from strand-β1 of the N-lobe and D13 forms a hydrogen bond with residue S116 of the hinge-loop as well as the backbone amino-group of R170 from the loop between strands β7 and β8 of the C-lobe (**Figure 4B**). The interaction partners of the YD-motif in medTK*b* (i.e. residues E43, S116 and R170) are conserved across the 462 TK sequences analysed in this study (99%, 99% and 96% sequence identity, respectively; including human TK and medTK*a* sequences). Attesting to the structural role of this motif, the experimental mutation of the conserved D residue to valine in human TK was shown to impair the interaction of the NL sequence with the kinase domain (Bogomolovas et al., 2021). Notably, the YD motif and its interacting residues are also conserved in TwcK homologues from invertebrates, showing the lowest RMSD values within the NL region when comparing available 3D-structures of TK and TwcK representatives (**Figure 4C**).

**Figure 4:**
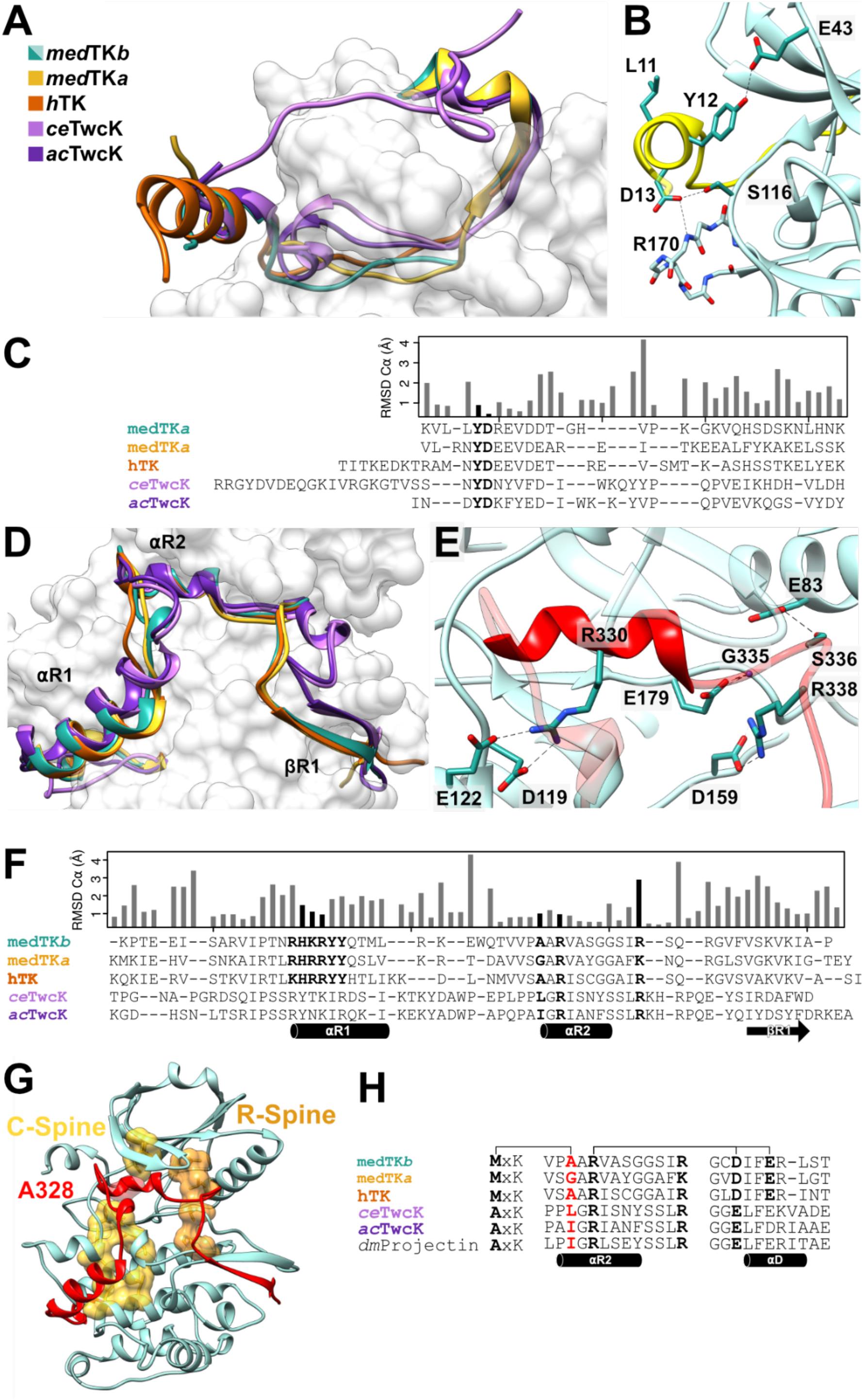
Comparison of N- and C-terminal tail extensions in titin(-like) kinases. **A.** N-terminal linker (NL) in available crystal structures of TK and TwcK homologues. Structures shown are excerpts from PDB codes: *ac*TwcK, 1KOB; *ce*TwcK, 3UTO; human TK, 6YGN. A 3D-model of medTK*a* (yellow) predicted using ColabFold (Mirdita et al., 2022) is included for completeness; **B.** Conserved interactions of the YD motif (residues 12-13) in the NL of medTK*b*; **C.** Structure-based sequence alignment of NL segments accompanied of a histogram of per-residue RMSD values for C⍺ atoms across all kinases shown in the alignment (calculated using UCSD chimera; https://www.cgl.ucsf.edu/chimera). The conserved YD motif is highlighted; **D.** C-terminal regulatory domain (CRD) of TK and TwcK homologues. Colouring as in A.; **E.** Interactions supported by helix ⍺R2 in medTK*b*. In medTK*b*, an additional (non-conserved) interaction is that of residue S336 with the conserved glutamate (E83) from the catalytic ⍺-helix C; **F.** Structure-based sequence alignment of CRD tails. Details as in C. The conserved [R/K]H[R/K]RYY and R-7x-R motifs are highlighted as well as the residue in position −2 of the R-7x-R motif that packs against strand β3; **G**. Structure of medTK*b* with its pre-assembled catalytic (C-, yellow) and regulatory (R-, orange) spines shown in surface representation. The C-spine completing residue A328 from helix ⍺R2 of the CRD is indicated; **H.** Sequence alignment of helix ⍺R2 in TK and TK-like sarcomeric kinases showing the R-7x-R motif and the interactions it supports (connector lines). The C-spine completing residue in position −2 is shown in red. The correlated variation of this residue with the ATP-binding AxK motif in strand β3 is evident. In TwcK, the E/D residue in the hinge region does not form the annotated interaction.

Functionally, data from *ce*TwcK showed that the interaction of the NL against the interlobular kinase hinge region contributes to the inhibition of phosphotransfer in this kinase (von Castelmur et al., 2012). In addition, based on force-biased molecular dynamics simulations, the interactions held by the YD motif have been proposed to act as a force-barrier that protects the kinase domain from stretch-unfolding induced by pulling forces in the sarcomere (Bogomolovas et al., 2021). The motif appears to confine stretch-unfolding to the NL tail and to prevent the propagation of stretch along the protein chain, thereby protecting the kinase domain from mechanical damage. Interestingly, the single nucleotide variant (SNV) D24728V affecting the YD motif of human TK has been found in patients with dilated cardiomyopathy, supporting the view that this motif might play an important functional role in titin (Bogomolovas et al., 2021). Other than the YD motif, sequences of the NL tail show little conservation (**Figure 4C**).

### The C-terminal CRD extension plays multiple inhibitory and stabilizing roles

The CRD tail extension of human TK binds tightly against the kinase domain blocking access to the active site (Mayans et al., 1998; Bogomolovas et al., 2014; Bogomolovas et al., 2021). It also contributes critically to kinase stability (Bogomolovas et al., 2022). Structurally, the CRD tail folds into two helices (the ⍺-helix ⍺R1 and the 3_10_-helix ⍺R2) and a C-terminal β-strand (strand βR1) (nomenclature as in Mayans et al., 1998) (**Figure 4D)**. Helix ⍺R1 is thought to be particularly required for TKs fold stability (Lange et al., 2005; Bogomolovas et al., 2022); this also being the case in other titin-like kinases such as PK1 from *Drosophila* obscurin (Zacharchenko et al., 2023). A feature of interest in helix ⍺R1 is the highly conserved motif [R/K]H[R/K]RYY (**Figure 4F**). The motif is partly solvent exposed on the surface of the helix, where it is freely accessible and makes a prime candidate for a binding interface. In human TK, the motif has been proposed to bind the autophagy receptor Nbr1 (Lange et al., 2005). However, subsequent *in vitro* studies could not confirm the interaction (Müller et al., 2006; Bogomolovas et al., 2021). Instead, data suggested that Nbr1 associates with ubiquitin-groups on TK once the kinase has undergone MuRF-mediated ubiquitination (Bogomolovas et al., 2021). Thus, whether this conserved motif supports interactions in TK is currently unclear. Notably, the motif is known to host an SNV (R32450W) that yields the sequence HWR (Lange et al., 2005). The potential association of this SNV with Hereditary Myopathy with Early Respiratory Failure (HMERF) is a matter of debate (Lange et al., 2014; Pfeffer & Chinnery, 2014).

Helix ⍺R2 and strand βR1 are the primary inhibitory elements of the CRD tail. Helix ⍺R2 is a short 3_10_-helix that penetrates the ATP-binding pocket, while strand βR1 forms a two-stranded antiparallel β-sheet with the activation loop that blocks the binding of any potential peptidic phosphorylation substrate. We observe that a conserved signature motif of this chain segment is RxxxxxxxR (here abbreviated as R-7x-R), where two arginine residues are separated by any 7 residues. The first R is in helix ⍺R2 and the second in the loop between helix ⍺R2 and strand βR1 (**Figure 4E,F**). In medTK*b,* the first arginine (R330) forms a salt bridge with residues D119 and D122 in the kinase hinge-loop and the second arginine (R338) forms a salt bridge with the catalytic aspartate (D159) (**Figure 4E**). The interactions held by the first arginine are conserved across all TK and TwcK homologues of known 3D structure. However, whether the second arginine interacts or not with the catalytic aspartate hinges on the presence or absence of the tyrosine rest in loop P+1 proposed to be part of the inhibitory mechanism of human TK (Mayans et al., 1998). In the absence of the tyrosine rest, the arginine interacts with the catalytic aspartate as observed in medTK*b* and the TwcKs from *C. elegans* and *Aplysia*. Specifically, in medTK*b* the catalytic aspartate D159 is embedded in a hydrogen bond network formed by residues R161, Q182 and R338 (**Figure 3D**). In the presence of the tyrosine residue, the catalytic aspartate does not interact with the arginine from the CRD, but with the tyrosine rest, as seen in various crystal structures of human TK (PDB: 1TKI, 4JNW, 6YGN) (**Figure 3D**). Thus, the catalytic aspartate appears to be part of an interaction switch where either the tyrosine residue from the P+1 loop or the arginine from the CRD are involved.

A further important feature of helix ⍺R2 is position −2 respect to the R-7x-R motif. By binding within the ATP-binding pocket, the residue at this position completes the catalytic C-spine of the kinase (**Figure 4G**). In canonical kinases, the C-spine is completed by the nitrogenated base of the bound ATP substrate. Thus, in TK and TwcK, the CRD tail mimics both ATP and peptide binding, completing the C-spine and maintaining the inhibited kinase in a primed catalytic conformation. In TwcK kinases, which are catalytically active, the C-spine is completed by a hydrophobic residue - commonly leucine or isoleucine - from helix ⍺R2 that packs against the small residue in position 1 of the <u>A</u>xK motif in strand β3. Both in medTK*b* and human TK, the motif in strand β3 unusually contains a bulky methionine residue, <u>M</u>xK, and position −2 in the CRD-helix ⍺R2 is a small residue, either alanine or glycine. It can be thus concluded that the residue size at this CRD position strongly anti-correlates with the size of the first residue in the AxK motif of strand β3, this co-variation resulting from a volume compensation of these mutually packing groups. Upon ATP binding, the residue from the CRD would be substituted by the intercalated insertion of the nitrogenated base moiety of the ATP. Our finding leads us to question whether the large ATP base could be well accommodated at this position; a deduction that adds to the previous speculation that the larger methionine in the MxK motif might lead to a non-productive binding mode of ATP in TK (Bogomolovas et al., 2014).

### The medTK*a* isoform contains a canonical DMG motif

To complete the analysis of TK kinases in medaka, we predicted the 3D-structure of the medTK*a* isoform using AlphaFold (**Figure S2A**) (Mirdita et al., 2022) and compare it to the experimental structures of human TK and medTK*b* (**Figure S2B**). The predicted model is in close agreement with the crystal structures of medTK*b* and human TK (RMSD_C*α*_=1.24 Å across all 344 aligned C_α_-atoms to medTK*b* and RMSD_C*α*_=0.89 Å across all 344 aligned C_α_-atoms to human TK [PDB: 6YGN], calculated using UCSD Chimera; https://www.cgl.ucsf.edu/chimera). The model of medTK*a* displays the same features as described in this work for medTK*b* and human TK. The hydrophobic spines are pre-formed, where V46, M59, I113, IVY158-160, M217 and L221 constitute the C-spine (M217 is equivalent to L224 in medTK*b*) and the residue in position −2 of helix ⍺R2 from the CRD-tail is a glycine (G320). The R-spine is composed of L80, L91, H150 and M173, where M173 is part of the canonical DFG-motif that here is DMG (ELG in medTK*b*). The DMG motif of medTK*a* contains the canonical aspartate and is also found in human CDK19 and CDK8, the latter having been shown to be an active kinase (Fryer et al., 2004). As human TK, medTK*a* has a tyrosine residue in the P+1 loop that is predicted to hydrogen bond to the catalytic aspartate (**Figure S2D**). The glycine-rich loop in medTK*a* contains a ARSQFG sequence that differs from the canonical GxGxxG motif, but that is not at odds with ATP-binding as this loop can accept a variety of residues (Matsunaga et al., 2024). In brief, medTK*a* shares the MxK atypical motif with human TK and medTK*b,* as well as sharing the presence of a potentially inhibitory tyrosine residue in loop P+1 with human TK. However, contrary to human TK and medTK*b*, it lacks the DtoE exchange in the DFG motif, having a canonical aspartate in this position. A future investigation of whether medTK*a* presents increased levels of phosphotransfer activity compared to medTK*b* and human TK is required to establish the functional significance of this observation.

### Comparison of sequence-motifs within and across PaSiMap clusters

Having explored sequence identifiers in crystal structures, we finally asked whether such features are representative of TK kinases at large or specific TK groups. To gain an insight into this question, we analysed sequence conservation in the identified PaSiMap clusters (**Figure 5**). We observed that the two deviating features of the kinase domain, i.e. the <u>M</u>xK and <u>E</u>xG, are present throughout all TK groups. Specifically, all 452 TK sequences from fish and mammals contained a large hydrophobic residue in the first position of the MxK motif. TK from mammals (n= 139) and TK isoforms *a* and *b* from neoteleostei (n=91 and n=100, respectively) contain a methionine residue at this position in 98.5%, 98.9% and 100% of its representatives. TK isoforms *a* and *b* from the older (and, therefore, more diversified) non-neoteleostei group (n= 60 and n=62, respectively) contain a methionine rest 40% and 77.4% of the times, respectively. Residues other than methionine across all groups were leucine (8.7%), isoleucine (0.6%) and phenylalanine (2.6%). It can be concluded that strand β3 of TK sequences hosts a θxK motif, where θ represents a bulky hydrophobic residue that is a general characteristic of TK kinases. Accordingly, position −2 in helix ⍺R2 of the CRD was consistently a small residue, most commonly an alanine across all groups (76.8% alanine, 15.2% serine/threonine, 4.8% glycine, 1.5% valine, only rarely 0.4% being a larger glutamate or glutamine residue). In contrast, the <u>E</u>xG motif presents a diversification that somewhat correlates with TK isoform. Mammalian TK and all isoform *b* (from both neoteleostei and non-neoteleostei) contain a glutamate residue (E) 100% of the time. TK isoform *a* largely contains a glutamate (E), but a number of representatives contain a canonical aspartate (D); namely, 17,6% and 6,7% occurrences in neoteleostei and non-neoteleostei, respectively (this amounts to 20 TK sequences, out of the 452 total in this study). Whether these TK*a* variants might deliver phosphotransfer catalysis or might have lost this capability through another mechanism is yet to be investigated. In that respect, it must be also noted that sequence variations in the ATP-binding GxGxxG loop also correlate with TK isoform. Mammalian TK and fish isoform *b* (from both neoteleostei and non-neoteleostei) have a conventional GxGxxG motif, while TK*a* variants from both fish groups have diversified loops. TK*a* from non-neoteleostei exhibit a RxGxxG motif while TK*a* from neoteleostei have an Ax[C/S]xxG sequence. In contrast to these features, rather modest conservation is found in the potentially inhibitory tyrosine from the P+1 loop. This residue is highly conserved only in mammals (96,4%), being modestly present in TK*a* isoforms from fish (28,6% in neoteleostei and 46,7% in non-neoteleostei) and totally absent from TK*b* isoforms. Thus, the functionality of this residue is not conserved in TK kinases.

**Figure 5.**
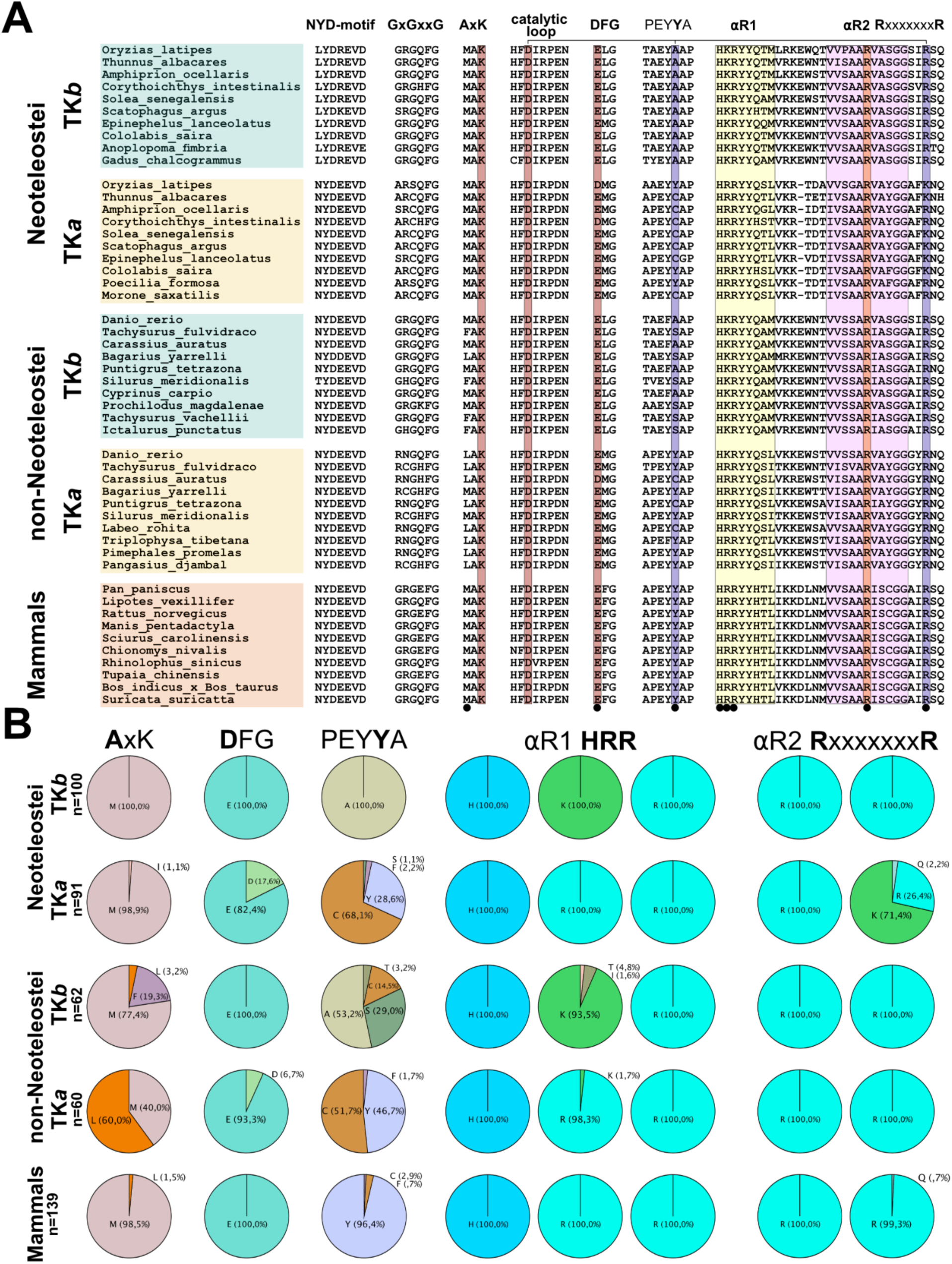
Comparison of conserved sequence motifs across PaSiMap TK clusters. Analysis of conservation in functional sequence motifs across the five clusters of TK sequences classed by PaSiMap (corresponding to TK isoforms *a* and *b* of neoteleostei, isoforms *a* and *b* of non-neoteleostei, and mammalian TK). **A.** Selected sequences for each group are shown. The sequence motifs are stated as headings in their canonical occurrence. The catalytic triad of kinases is highlighted in brown, these are the lysine residue in the AxK motif, the aspartate residue in the catalytic loop and the D/E rest of the DFG-motif. The two different modes of inhibition (tyrosine in P+1 loop or arginine from the CRD) of the catalytic aspartate are marked in blue and connected by a horizontal bar on top of the sequence. The helix ⍺R1 of the CRD is highlighted in yellow. **B.** Quantitation of conserved residues across the sequence groups is shown and given in percentages. The selected residue positions quantitated in the chart correspond to the residue in bold in the sequence above each chart. The same residues are also indicated with small black circles in the multiple sequence alignment in A.

Remarkably, a very high conservation of specific features in the NL and CRD flanking tails is also observed. In the NL segment, the [N/L]YD motif is strictly present in all TK sequences here studied, irrespective of animal group. In the CRD, the almost strict conservation of the [K/R]H[<u>R/K]R</u>YY motif in helix ⍺R1 of all TK sequences is also striking, especially as the two central positively charged residues (99% K/R and 100% R, underlined) are exposed to solvent and facing away from the kinase domain. The conservation of the R-7x-R motif in helix ⍺ R2 is also striking and can be explained by the role that this motif plays in the packing of the tail against the kinase domain.

## Conclusion

Sarcomeric kinases are a group of kinases and pseudokinases thought to play mechanosensory roles in muscle. The best studied representatives of this group are the close homologues human TK and TwcK from *C. elegans*. TwcK is an active kinase that undergoes autophosphorylation (Williams et al., 2018) and has demonstrated activity on model substrates (Heierhorst et al., 1996; von Castelmur et al., 2012). The ablation of TwcK’s phosphotransfer activity through the inactivating point mutation KtoA has a phenotypic outcome in which *C. elegans* worms swim at an accelerated pace yet display reduced overall fitness (Matsunaga et al., 2017). In contrast, human TK is a pseudokinase and while its scaffolding functions are well established (Lange et al., 2005; Bogomolovas et al., 2021), its phosphotransfer activity is thought to be ablated or highly reduced (Bogomolovas et al., 2014; Bogomolovas et al., 2022). Accordingly, no validated substrate has been established for TK to date (Bogomolovas et al., 2022). Here, we set out to examine sequence features in TK across fish and mammals that cover long evolutionary distances, aiming to reveal if the pseudokinase character of TK is conserved through evolution.

Our study reveals that, despite sequence variations, TK domains are strongly conserved structurally, including their flanking tails, and that their pseudokinase features are preserved through evolution. In particular, the θxK motif (where θ is a bulky, hydrophobic residue) and the EFG motif, both thought to hinder phosphotransfer activity, are largely conserved throughout TK sequences. Our elucidation of the crystal structure of medTK*b* leads us to rationalize the conservation of the θ residue in the first motif as a covariation with the presence of a small residue in the CRD sequence that penetrates the ATP pocket and packs against it, adding to the notion that TK cannot bind ATP in a canonical fashion. Using a previously reported multiple sequence alignment of human kinases (Modi & Dunbrack, 2019) we identified that RIPK1 (Chan et al., 2003), NEK8 (Holland et al., 2002) and Eif2ak4 (Sood et al., 2000) kinases contain isoleucines in position θ and LMTK2 (Wang & Brautigan, 2006) contains leucine, yet all being active kinases. Similarly, the ATR-kinase contains a methionine as θ residue and is also proposed to be active (Kim et al., 1999). On the other hand, RYK kinase contains a phenylalanine residue in this position and is known to be catalytically inactive (Hovens et al., 1992), not detectably binding nucleotides or cations (Murphy et al., 2014). The extent to which the methionine residue in the MxK motif of TK affects its phosphotransfer catalysis awaits future clarification.

The reasons for the strict conservation of the glutamate residue in the EFG motif are, however, unclear. Although the glutamate side chain reaches to the backbone of the CRD tail and binds a conserved glycine residue (in *ce*TwcK the canonical aspartate interacts with water molecules in the CRD-bound state), this intrachain interaction alone does not explain conservation. As evolutionarily only those protein residues with functional implications need to be retained, lacking the constraint of performing phosphotransfer allows pseudokinases to diverge in catalytically critical residues (Murphy et al., 2014; Kwon et al., 2019). Notably, if the physiological functionality of TK did not require it to display phosphotransfer activity and given the absence of structural constraints, a variety of inactivating residues could be expected to occur in lieu of E/D at this position. The high conservation of the glutamate residue in this motif and its chemical resemblance to the canonical aspartate lead us to speculate that this might be a compensatory residue exchange, possibly correlated to the θxK motif, and directed to enable certain phosphotransfer activity. In this regard, it is interesting to note that our analysis has identified a set of 20 TK sequences (including medTK*a*) containing the canonical DFG motif. All 20 kinases have the θxK motif (16 sequences have M, 4 sequences L). A future characterization of these TK variants will be highly informative to further elucidate the phosphotransfer properties of TK in the muscle sarcomere.

Beyond the kinase domain, the NL and CRD extensions of TKs show remarkable structural similarity even though their sequence conservation is modest. Conservation is primarily limited to four motifs: (i) the YD-motif of the NL; (ii) a [R/K]H[R/K]RYY motif in helix ⍺ R1 of the CRD; (iii) a R-7x-R motif in the CRD helix ⍺ R2; and (iv) a small residue in position −2 of the R-7x-R motif. With the exception of motif [R/K]H[R/K]RYY (ii), whose function is unknown, the role of the conserved motifs (i, iii, iv) is to mediate the packing of the tails against the kinase domain. The YD-motif of the NL is the primary anchoring point of the NL to the kinase domain, its mutation disrupting the interaction (Bogomolovas et al., 2021). The R-7x-R motif and its −2 position anchor helix ⍺ R2 from the CRD onto the kinase and into the ATP binding pocket. In TK, residue θ is a bulky hydrophobic and position −2 a small residue, together they complete the C-spine. In TwcK from invertebrates, residue θ is small (often an alanine), while position −2 houses a larger hydrophobic residue that completes the C-spine. We conclude that there are structural constraints on the residue in position −2 and that its divergence is part of the pseudokinase sequence signature of TK.

Finally, we identify a clear segregation of TK isoforms in fish, evidenced by sequence clustering (**Figure 1A,C)**. The segregation of TK sequences resembles that of full-length chain sequences in a selected set chosen for analysis (**Figure S1**). This leads us to propose that TK domains can be utilized as a proxy in the classification of titin genes from fish. Specifically, adopting the titin gene nomenclature used for zebrafish, we propose a classification of currently non-annotated titin genes into *a* and *b* isoforms (*ttna* and *ttnb* genes) that can assist future comparative studies. Thus, a curated gene annotation is provided here as supplementary material (**Supp Mat Table S1**). Finally, the molecular analysis presented here will contribute to further our understanding of titin properties across mammalian, zebrafish and medaka *in vivo* model systems, advancing our interpretation of physiological data derived from their study.

## Supporting information

Supplementary Interactive PasiMap 3D-vector map S2

Supplementary Interactive PasiMap 3D-vector map S3

## Data availability

Structure coordinates and experimental diffraction data have been deposited with the PDB (entry 9QJ2). X-ray diffraction images have been deposited with Zenodo (http://doi.org/10.5281/zenodo.14922781)

## Acknowledgments

We gratefully acknowledge financial support from KoRS-CB and AFF from the University of Konstanz and the provision of synchrotron radiation time by EMBL Hamburg at the PETRA III storage ring (DESY, Hamburg, Germany).

## Author contribution

TD and OM conceived the study; PG performed cloning, protein production, and crystallization of medTK*b*; TD and OM performed crystallographic structural elucidation, TD performed bioinformatics analysis; TD and OM wrote the manuscript.

## Supplementary Materials

### SuppMat_TableS1-GeneNames.xlsx

List of protein entries used in the sequence comparison in this study. Entries are grouped according to their PaSiMap cluster classification, where groups correspond to mammals, neoteleostei TK isoform a (TK*a*), non-neoteleostei TK*a*, neoteleostei TK isoform b (TK*b*) and non-neoteleostei TK*b.* For each column the NCBI database access code to the gene is given, followed by the residue range corresponding to the TK sequence analyzed, the gene name and the source organism.

### SuppMat_S2 (3DPaSiMap_Fish_TK.html): Interactive PaSiMap vector map revealing similarity clusters of TK sequences from fish

Shown are dimensions 2, 3 and 4 of the PaSiMap cluster map of 322 TK sequences from fish. Circles and empty diamonds indicate sequences from neoteleostei and non-neoteleostei, respectively. Yellow corresponds to TK isoform a and teal to TK isoform b, as named in this manuscript. A black cross indicates the origin of coordinates. (Plot created in R-Studio; http://www.rstudio.com/).

### SuppMat_S3 (3DPaSiMap_Fish_Mammals_TK.html): Interactive map of PaSiMap clustering TK sequences from fish and mammals

Shown are dimensions 2, 3 and 4 of the PaSiMap vector map of 461 TK sequences from fish and mammals. Mammalian TK are brown squares, circles and diamonds indicate sequences from neoteleostei and non-neoteleostei, respectively. Yellow corresponds to TK isoform a and teal to TK isoform b, as named in this manuscript. A black cross indicates the origin of coordinates. (Plot created in R-Studio; http://www.rstudio.com/).

**Figure S1:**
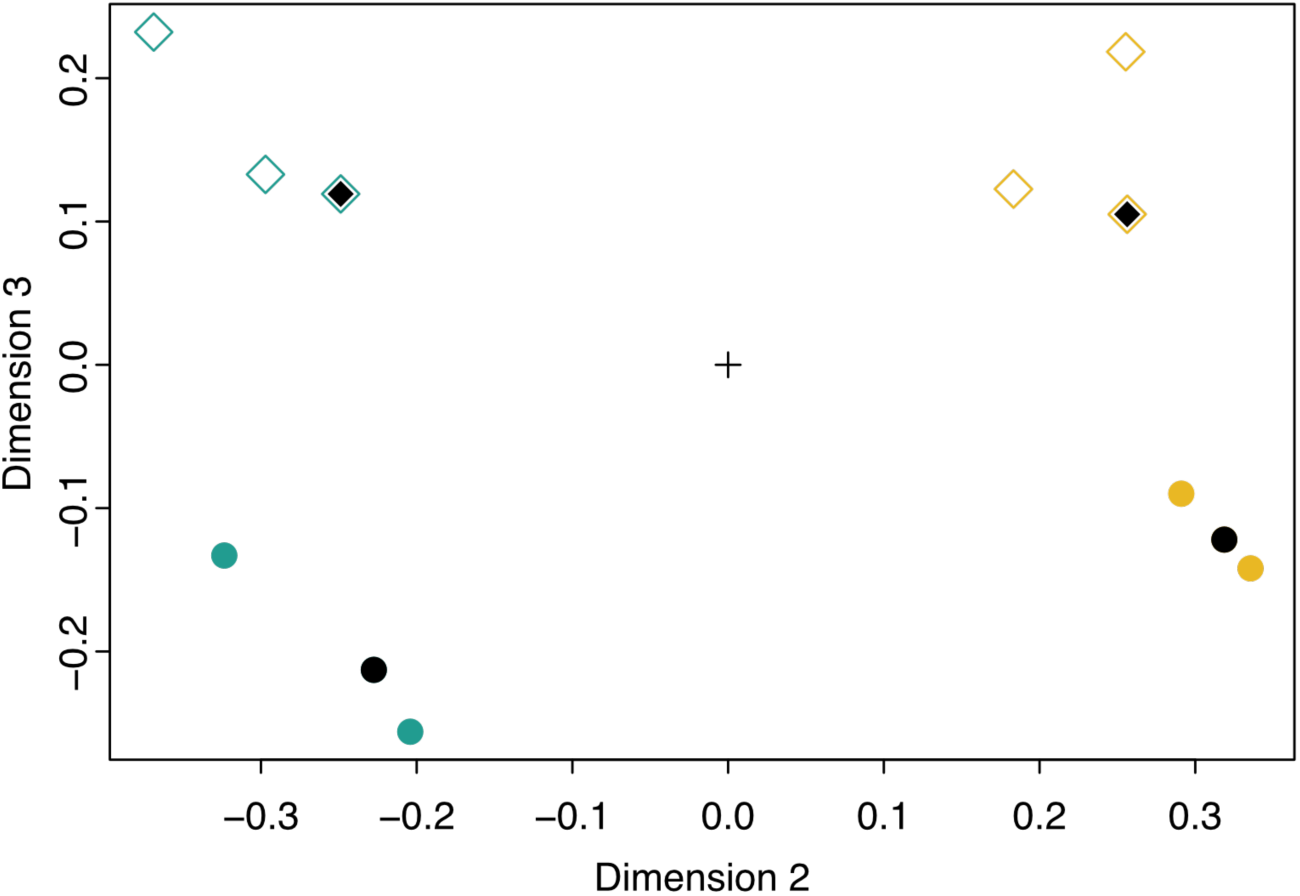
PaSiMap vector map of full-length titin protein sequences from selected fish representatives. PaSiMap analysis of full-length titin isoforms from the neoteleostei *Amphiprion ocellaris, Oryzias latipes* and *Thunnus albacares*; and non-neoteleostei *Danio rerio*, *Tachysurus fulvidraco* and *Carassius auratus*. Fish groupings are indicated by rhombic and circular data points, where rhomboids are used for non-neoteleostei and circles indicate neoteleostei. The model organisms representative of each group, Zebrafish (*Danio rerio*; ◆) and medaka (*Oryzias latipes*; □) are indicated with black symbols. Yellow and teal colours indicate isoforms *a* and *b* as deduced from the clustering of titin kinase sequences in Fig 1A. The result shows that the clustering of full-length titin sequences reproduces the clustering of titin kinase sequences, so that the TK domain is a valid surrogate for the classification of full-length titin protein isoforms. Based on this analysis, we conclude that entries corresponding to titin isoform a are: *Amphiprion ocellaris* (XP_054871008.1), *Oryzias latipes* (XP_023806459.1), *Thunnus albacares* (XP_044221301.1), *Danio rerio* (ABG48500.1), *Tachysurus fulvidraco* (XP_047670400.1), *Carassius auratus* (XP_026074665.1); isoforms b are *Amphiprion ocellaris* (XP_054871216.1), *Oryzias latipes* (XP_023806503.1), *Thunnus albacares* (XP_044222812.1), *Danio rerio* (ABG48499.1), *Tachysurus fulvidraco* (XP_047670412.1) and *Carassius auratus* (XP_026127101.1).

**Figure S2:**
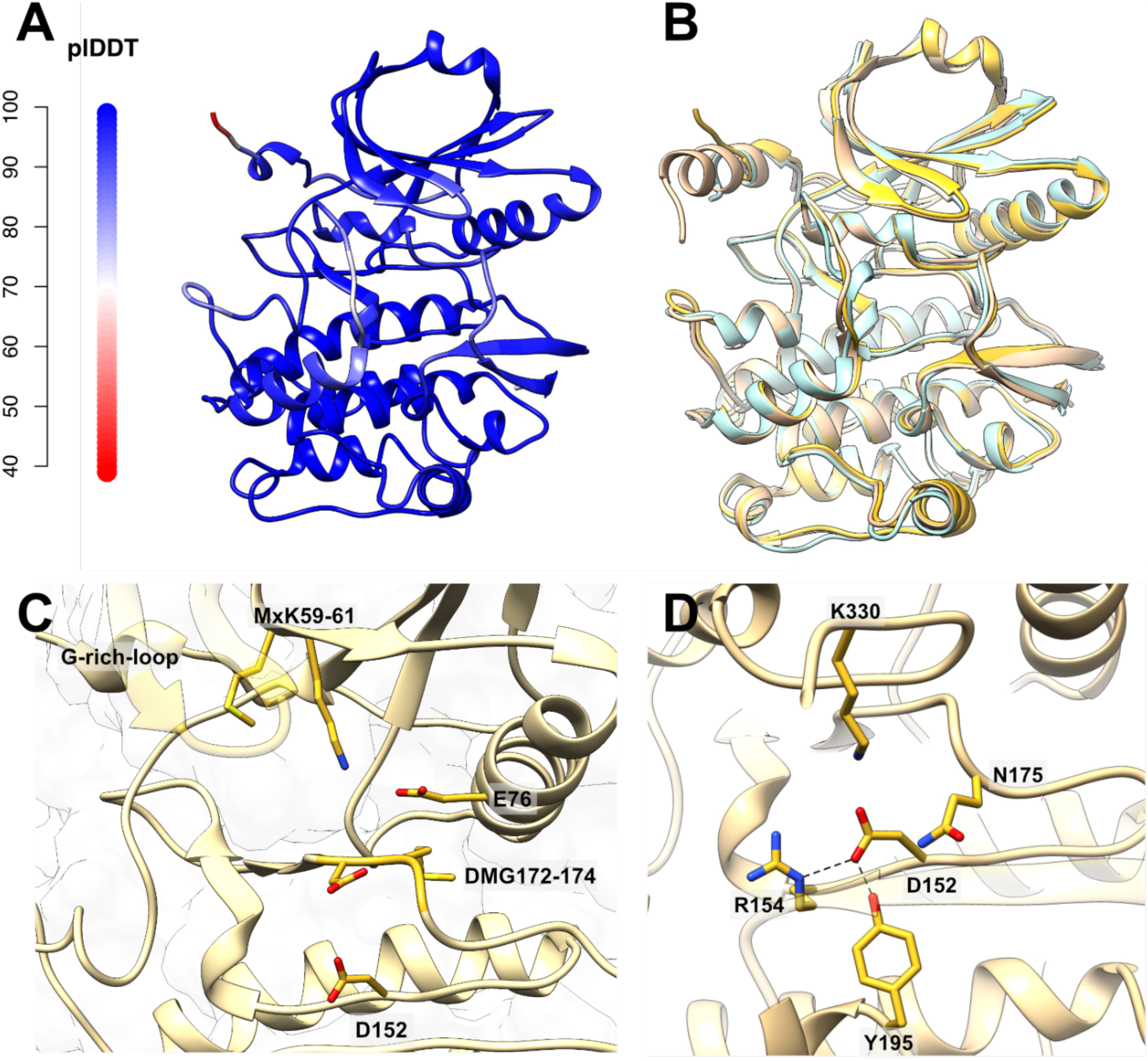
AlphaFold prediction of the 3D-model of *med*TK*a*. **A.** 3D-model of *med*TK*a* predicted using AlphaFold2 (https://colab.research.google.com/github/sokrypton/ColabFold/blob/main/AlphaFold2.ipynb; Mirdita et al., 2022) (NCBI XP_023806459.1; titin isoform X10; residues 29360-29710). The structure is coloured according to prediction confidence, where the colour gradient corresponds to the values of the pLDDT score as indicated by the scale bar on the left. Regions with a pLDDT below 70 should be treated with caution as the modelling is deemed unreliable then; **B.** Superimposition of the model predicted for *med*TK*a* (yellow) with the crystal structures of *med*TK*b* (light blue) and human TK (PDB: 6YGN; beige). RMSD_C*α*_=1.24 Å across all 344 aligned C_α_-atoms to medTK*b* and RMSD_C*α*_=0.89 Å across all 344 aligned C_α_-atoms to human TK, calculated using UCSD Chimera; https://www.cgl.ucsf.edu/chimera); **C.** Functional groups of the predicted medTK*a*. The CRD tail is removed for visual clarity; **D** Interactions sustained by the catalytic aspartate in the predicted 3D-model *med*TK*a*. Hydrogen bonds are shown as dotted lines.

## References

Bang ML, Centner T, Fornoff F, Geach AJ, Gotthardt M, McNabb M, Witt CC, Labeit D, Gregorio CC, Granzier H, Labeit S. (2001). The complete gene sequence of titin, expression of an unusual approximately 700-kDa titin isoform, and its interaction with obscurin identify a novel Z-line to I-band linking system. Circulation Research, 89(11):1065–1072.

Bogomolovas J, Fleming JR, Franke B, Manso B, Simon B, Gasch A, Markovic M, Brunner T, Knöll R, Chen J, Labeit S, Scheffner M, Peter C, Mayans O. (2021). Titin kinase ubiquitination aligns autophagy receptors with mechanical signals in the sarcomere. EMBO Reports, 22(10):e48018.

Bogomolovas J, Gasch A, Simkovic F, Rigden DJ, Labeit S, Mayans O. (2014). Titin kinase is an inactive pseudokinase scaffold that supports MuRF1 recruitment to the sarcomeric M-line. Open Biol., 4(5):140041.

Bogomolovas J, Gravenhorst P, Mayans O. (2022). Chapter Seven - Production and analysis of titin kinase: Exploiting active/inactive kinase homologs in pseudokinase validation. In N. Jura & J. M. Murphy (Eds.), Meth. Enzymol. (Vol. 667, pp. 147–181). Academic Press.

Brown EJ, Beal PA, Keith CT, Chen J, Shin TB, Schreiber SL. (1995). Control of p70 s6 kinase by kinase activity of FRAP in vivo. Nature, 377(6548):441–446.

Chan FK-M, Shisler J, Bixby JG, Felices M, Zheng L, Appel M, Orenstein J, Moss B, Lenardo MJ. (2003). A Role for Tumor Necrosis Factor Receptor-2 and Receptor-interacting Protein in Programmed Necrosis and Antiviral Responses. J. Biol. Chem., 278(51):51613–51621.

de Diego I, Kuper J, Bakalova N, Kursula P, Wilmanns M. (2010). Molecular basis of the death-associated protein kinase-calcium/calmodulin regulator complex. Science Signaling, 3(106):ra6.

Emsley P, Lohkamp B, Scott WG, Cowtan K. (2010). Features and development of Coot. Acta Cryst. Section D, Biological Crystallography, 66(Pt 4):486–501.

Endicott JA, Noble MEM, Johnson LN. (2012). The structural basis for control of eukaryotic protein kinases. Ann. Rev. Biochem., 81:587–613.

Fryer CJ, White JB, Jones KA. (2004). Mastermind recruits CycC:CDK8 to phosphorylate the Notch ICD and coordinate activation with turnover. Mol. Cell, 16(4):509–520.

Gräter F, Shen J, Jiang H, Gautel M, Grubmüller H. (2005). Mechanically Induced Titin Kinase Activation Studied by Force-Probe Molecular Dynamics Simulations. Biophys. J., 88(2):790–804.

Heierhorst J, Tang X, Lei J, Probst WC, Weiss KR, Kemp BE, Benian GM. (1996). Substrate Specificity and Inhibitor Sensitivity of Ca2+/S100-dependent Twitchin Kinases. Europ. J. Biochem., 242(3):454–459.

Holland PM, Milne A, Garka K, Johnson RS, Willis C, Sims JE, Rauch CT, Bird TA, Virca GD. (2002). Purification, Cloning, and Characterization of Nek8, a Novel NIMA-related Kinase, and Its Candidate Substrate Bicd2. J. Biol. Chem., 277(18):16229–16240.

Hovens CM, Stacker SA, Andres AC, Harpur AG, Ziemiecki A, Wilks AF. (1992). RYK, a receptor tyrosine kinase-related molecule with unusual kinase domain motifs. Proc. Nat. Acad. Sc. USA, 89(24):11818–11822.

Hu S-H, Parker MW, Yi Lei J, Wilce MCJ, Benian GM, Kemp BE. (1994). Insights into autoregulation from the crystal structure of twitchin kinase. Nature, 369(6481):581– 584.

Kabsch W. (2010). XDS. Acta Crysta Section D: Biological Crystallography, 66(2):2.

Kim S-T, Lim D-S, Canman CE, Kastan MB. (1999). Substrate Specificities and Identification of Putative Substrates of ATM Kinase Family Members. J. Biol. Chem., 274(53):37538–37543.

Kobe B, Heierhorst J, Feil SC, Parker MW, Benian GM, Weiss KR, Kemp BE. (1996). Giant protein kinases: Domain interactions and structural basis of autoregulation. EMBO J, 15(24):6810–6821.

Kornev AP, Taylor SS. (2015). Dynamics driven allostery in protein kinases. Trends in Biochem. Sc., 40(11):628–647.

Krüger M, Kötter S. (2016). Titin, a Central Mediator for Hypertrophic Signaling, Exercise-Induced Mechanosignaling and Skeletal Muscle Remodeling. Front. Physiol, 7:76.

Kwon A, Scott S, Taujale R, Yeung W, Kochut KJ, Eyers PA, Kannan N. (2019). Tracing the origin and evolution of pseudokinases across the tree of life. Sci. Signaling, 12(578): eaav3810.

Labeit S, Kolmerer B. (1995). Titins: Giant proteins in charge of muscle ultrastructure and elasticity. Science, 270(5234):293–296.

Lange S, Edström L, Udd B, Gautel M. (2014). Reply: Hereditary myopathy with early respiratory failure is caused by mutations in the titin FN3 119 domain. Brain, 137(Pt 6):e279.

Lange S, Xiang F, Yakovenko A, et al. (2005). The kinase domain of titin controls muscle gene expression and protein turnover. Science, 308(5728):1599–1603.

Langer G, Cohen SX, Lamzin VS, Perrakis A. (2008). Automated macromolecular model building for X-ray crystallography using ARP/wARP version 7. Nature Protocols, 3(7):1171–1179.

Liebschner D, Afonine PV, Baker ML, et al. (2019). Macromolecular structure determination using X-rays, neutrons and electrons: Recent developments in Phenix. Acta Cryst. Section D: Structural Biology, 75(10):10.

Linke WA. (2023). Stretching the story of titin and muscle function. Journal of Biomechanics, 152:111553.

Mace PD, Wallez Y, Egger MF, Dobaczewska MK, Robinson H, Pasquale EB, Riedl SJ. (2013). Structure of ERK2 bound to PEA-15 reveals a mechanism for rapid release of activated MAPK. Nature Comm., 4(1):1681.

Manning G, Whyte DB, Martinez R, Hunter T, Sudarsanam S. (2002). The protein kinase complement of the human genome. Science, 298(5600):1912–1934.

Matsunaga Y, Hwang H, Franke B, Williams R, Penley M, Qadota H, Yi H, Morran LT, Lu H, Mayans O, Benian GM. (2017). Twitchin kinase inhibits muscle activity. Mol. Biol. Cell, 28(12):1591–1600.

Matsunaga Y, Qadota H, Ghazal N, Lesanpezeshki L, Dorendorf T, Moody JC, Ahier A, Matheny CJ, Vanapalli SA, Zuryn S, Mayans O, Kwong JQ, Benian GM. (2024). Protein kinase 2 of the giant sarcomeric protein UNC-89 regulates mitochondrial morphology and function. Comm. Biol., 7(1):1342.

Mayans O, van der Ven PFM, Wilm M, Mues A, Young P, Fürst DO, Wilmanns M, Gautel M. (1998). Structural basis for activation of the titin kinase domain during myofibrillogenesis. Nature, 395(6705):863–869.

McCoy AJ, Grosse-Kunstleve RW, Adams PD, Winn MD, Storoni LC, Read RJ. (2007). Phaser crystallographic software. J. Appl. Cryst., 40(Pt 4):658–674.

Modi V, Dunbrack RL. (2019). A Structurally-Validated Multiple Sequence Alignment of 497 Human Protein Kinase Domains. Sc. Rep., 9(1):Article 1.

Mirdita M, Schütze K, Moriwaki Y, Heo L, Ovchinnikov S, Steinegger M. (2022). ColabFold: Making protein folding accessible to all. Nature Methods, 19(6):679–682.

Morell T, Procter J, Barton GJ, Diederichs K, Mayans O, Fleming JR. (2024). Protocol for sequence clustering with PaSiMap in Jalview. STAR Protoc. 6(1):103603.

Müller S, Kursula I, Zou P, Wilmanns M. (2006). Crystal structure of the PB1 domain of NBR1. FEBS Lett. 580(1):341–4.

Murphy JM, Zhang Q, Young SN, et al. (2014). A robust methodology to subclassify pseudokinases based on their nucleotide-binding properties. Biochem J, 457(2): 323– 334.

Parey E, Louis A, Montfort J, Guiguen Y, Crollius HR, Berthelot C. (2022). An atlas of fish genome evolution reveals delayed rediploidization following the teleost whole-genome duplication. Genome Research, 32(9), 1685–1697.

Pfeffer G, Chinnery PF. (2014). Reply: Hereditary myopathy with early respiratory failure is caused by mutations in the titin FN3 119 domain. Brain, 137(6):e280.

Porto D, Matsunaga Y, Franke B, Williams RM, Qadota H, Mayans O, Benian GM, Lu H. (2021). Conformational changes in twitchin kinase in vivo revealed by FRET imaging of freely moving C. elegans. eLife, 10:e66862.

Puchner EM, Alexandrovich A, Kho AL, Hensen U, Schäfer LV, Brandmeier B, Gräter F, Grubmüller H, Gaub HE, Gautel M. (2008). Mechanoenzymatics of titin kinase. Proc. Nat. Acad. Sc. USA 105(36):13385–13390.

Qi M, Clark J, Moody ERR, Pisani D, Donoghue PCJ. (2024). Molecular Dating of the Teleost Whole Genome Duplication (3R) Is Compatible With the Expectations of Delayed Rediploidization. Genome Biol. Evol., 16(7):evae128.

Schu PV, Takegawa K, Fry MJ, Stack JH, Waterfield MD, Emr SD. (1993). Phosphatidylinositol 3-kinase encoded by yeast VPS34 gene essential for protein sorting. Science, 260(5104):88–91.

Seeley M, Huang W, Chen Z, Wolff WO, Lin X, Xu X. (2007). Depletion of zebrafish titin reduces cardiac contractility by disrupting the assembly of Z-discs and A-bands. Circulation Res., 100(2):238–245.

Sood, R., Porter, A. C., Olsen, D., Cavener, D. R., & Wek, R. C. (2000). A Mammalian Homologue of GCN2 Protein Kinase Important for Translational Control by Phosphorylation of Eukaryotic Initiation Factor-2α. Genetics, 154(2), 787–801.

Su K, Mayans O, Diederichs K, Fleming JR. (2022). Pairwise sequence similarity mapping with PaSiMap: Reclassification of immunoglobulin domains from titin as case study. Comp. Struct. Biotech. J., 20:5409–5419.

Taylor SS, Kornev AP. (2011). Protein kinases: Evolution of dynamic regulatory proteins. Trends in Biochem. Sci., 36(2):65–77.

Temmerman K, Simon B, Wilmanns M. (2013). Structural and functional diversity in the activity and regulation of DAPK-related protein kinases. FEBS J., 280(21):5533–5550.

von Castelmur E, Strumpfer J, Franke B, Bogomolovas J, Barbieri S, Qadota H, Konarev PV, Svergun DI, Labeit S, Benian GM, Schulten K, Mayans O. (2012). Identification of an N-terminal inhibitory extension as the primary mechanosensory regulator of twitchin kinase. Proc. Nat. Acad. Sc. USA, 109(34):13608–13613.

Wang H, Brautigan DL. (2006). Peptide Microarray Analysis of Substrate Specificity of the Transmembrane Ser/Thr Kinase KPI-2 Reveals Reactivity with Cystic Fibrosis Transmembrane Conductance Regulator and Phosphorylases. Mol. Cell. Proteomics, 5(11):2124–2130.

Williams RM, Franke B, Wilkinson M, Fleming JR, Rigden DJ, Benian GM, Eyers PA, Mayans O. (2018). Autophosphorylation Is a Mechanism of Inhibition in Twitchin Kinase. J Mol Biol, 430:793–805.

Wu J, Tseng YD, Xu C-F, Neubert TA, White MF, Hubbard SR. (2008). Structural and biochemical characterization of the KRLB region in insulin receptor substrate-2. Nat. Struct. Mol. Biol., 15(3):251–258.

Zacharchenko T, Dorendorf T, Locker N, Van Dijk E, Katzemich A, Diederichs K, Bullard B, Mayans O. (2023). PK1 from Drosophila obscurin is an inactive pseudokinase with scaffolding properties. Open Biol., 13(4):220350.

Zou J, Tran D, Baalbaki M, et al. (2015). An internal promoter underlies the difference in disease severity between N- and C-terminal truncation mutations of Titin in zebrafish. eLife, 4:e09406.

